# Platelet proteolytic machinery assessment in Alzheimer’s Diseases

**DOI:** 10.1101/760116

**Authors:** Roy G. Muriu, Jessica M. Sage, Abdulbaki Agbas

**Affiliations:** Department of Basic Sciences, Kansas City University of Medicine and Biosciences, Kansas City, MO 64106, USA

**Keywords:** Autophagy, Proteasome, platelets, Alzheimer’s disease, protein aggregation, TDP-43, neurodegeneration

## Abstract

**Aim:** Platelets provide substantial information about the proteolytic system profile in neurodegenerative diseases. Assessment of autophagy and proteasome target proteins in platelets may reflect tissue proteolytic machinery profile in central nervous system in Alzheimer’s diseases (AD). We aimed to demonstrate the optimum assay conditions and identify target proteins in platelet proteolytic machinery.

**Methods:** Platelet samples were obtained from clinically verified AD patients and age-matched non-demented control subjects that were recruited by University of Kansas Alzheimer’s disease Center.

Autophagosome participating proteins in platelets were identified by Western blotting analysis. Standard gel electrophoresis and electro transfer apparatus were used for protein transfer onto the membrane. Several antibodies were tested to identify the best working antibodies, and their concentrations were optimized. An ELISA kit was used for platelet proteasome protein determination. Infrared imaging technology was used for visualizing the proteins on the membrane.

**Results:** Autophagosome participating proteins showed elevated levels in AD patient platelet cytosol. Only LC3-I autophagosome protein levels were significantly elevated. The concentrations of platelet lysate proteasome were assessed. AD patient’s proteasome levels were elevated but they were statistically not important as compared to controls.

**Conclusions:** Platelets can be used for assessing whether proteolytic system is functional. Blood-based sampling from human donors is less-invasive and analyzing platelet proteolytic system profile may help to develop pharmaceutical intervention approaches for neurodegenerative diseases in general.

## INTRODUCTION

The purpose of this study is to demonstrate that human blood-derived platelets may provide critical information about malfunctioned proteolytic machinery leading to diseased protein aggregation in the neurodegenerative disease. Alzheimer’s disease (AD) is mainly characterized by protein aggregations and deposition that are toxic and lethal to cellular structures in the central nervous system ^[1]^. The proteolytic system that includes autophagy and proteasome pathways, degrade and allow recyclization of targeted molecules and organelles in all eukaryotic cells (Fig.1). The reason certain proteins are allowed to aggregate is the inhibition of proteolysis and down regulation of effector proteins that stimulate the pathogenesis of the degradation ^[2]^. A dysfunctional proteolytic system, including autophagy and proteasome, is implicated in the pathogenesis of AD. Autophagy is a lysosomal degradative process by which cellular residents are inducted into homeostasis through quality control by clearance of pathogenic proteins, recycling of macromolecules, and response to energy requirements ^[3]^. In neurodegeneration, this system has been proven dysfunctional and therefore associated with lack of proper protein disposal and thereby aggregation ^[4]^. A two-stage autophagy impairment (i.e., induction and lysosomal acidification) leads to pathogenesis of AD ^[2]^. During the induction process, autophagy requires the release and presence of beclin-1 protein from endoplasmic reticulum and formation of a multimeric complex ^[2]^.This is the site of autophagy initiation through vesicle nucleation then formation of isolation membrane ^[5]^. Beclin-1 is a crucial regulator of autophagy. Its expression in the hippocampus is decreased at the RNA and protein level in AD with advanced age. This protein is necessary for nucleation of a phagophore membrane before the autophagosome vesicle is fully formed ^[6]^. This concept produced comparative results against control subjects that demonstrated an increase in protein aggregation ^[7]^.

**Figure-1.**
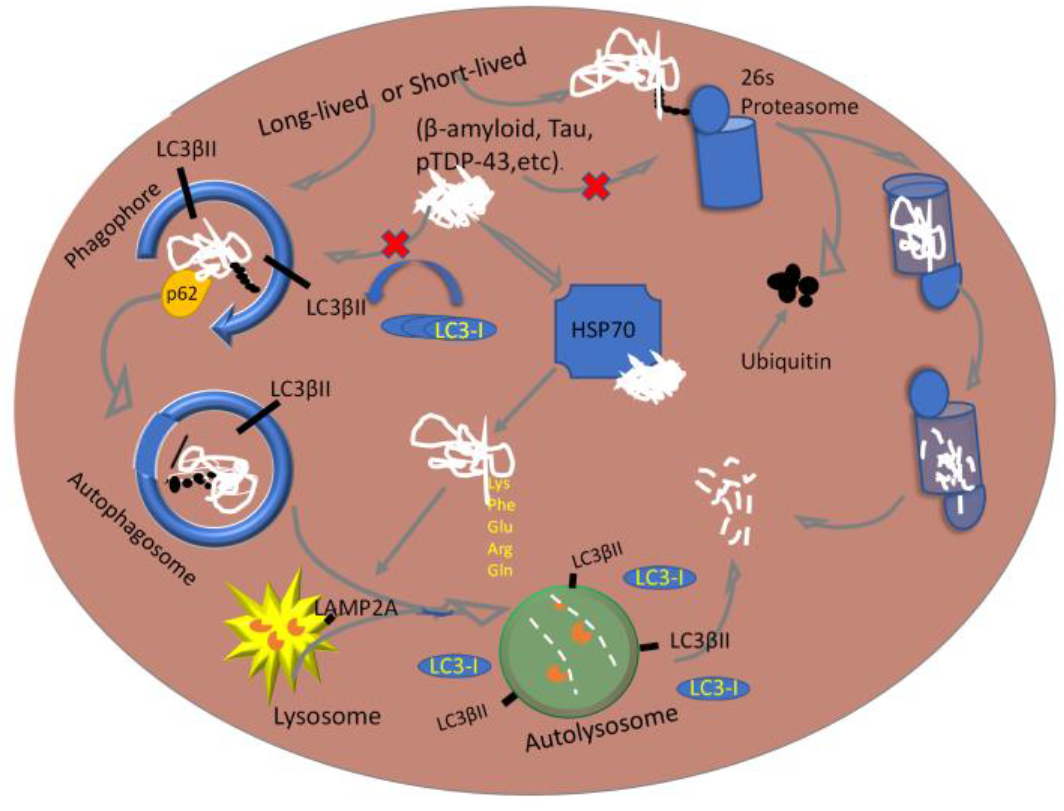
Schematic diagram of the proteolytic system in autophagic and proteasomal pathways.

Proteasomes are molecular machines that degrade aberrant proteins through polyubiquitination process followed by recognition by regulatory particle, deubiquitylation by deubiquitinase (DUB), and finally degradation of proteins into fragments ^[8]^. In AD, the components that are linked to this pathology are abnormal formation of ubiquitin and activity inhibition ^[9, 10]^. For example, paired helical filaments of tau (PHF-tau) bind to proteasomes and thereby reduce its activity ^[9]^, resulting less polyubiquitination Without this step, a protein without proper ubiquitination cannot be recognized by the regulatory particle of the proteasomes ^[11]^. Therefore, the autophagy and proteasome activities are a critical component and can be measured in platelets. Although the functions of autophagy within platelets are largely unclear but thus far, it is known that, its impairment leads to a lack of platelet aggregation and adhesion ^[12]^.

We have demonstrated that a TAR-DNA/RNA binding protein (TDP-43) and its phosphorylated derivative (pTDP-43) levels were elevated in platelets obtained from AD patients as part of the blood-based biomarker development studies ^[13]^. In this pilot study, we assessed the profile of target proteins for proteolytic machinery in platelet lysates obtained from AD patients and non-demented control subjects.

## METHODS

1. *Human Platelets* : AD patients and age-matched non-demented control subject human platelets were obtained from the biorepository of University of Kansas Alzheimer’s Disease Center under the approved IRB protocol (KUADC#11132). 5-6 days old platelets were obtained from a local community blood center (CBC) for initial studies before analyzing AD patients and control human platelet samples.
2. *Materials*:

2.1. Antibodies:

2.1.1. MBL antibodies (MBL International, 15A Constitution way, Woburn, MA 01801, USA): Anti-LC3(pAb, PM036Y), Anti-LC3 (8E10, mAb, M186-3Y), Anti-LC3 (4E12, mAb, M152-3Y), Anti-Beclin-1 (pAb, PD017Y), Anti-Atg16L (pAb, PM040Y), Anti-p62 (SQSTM1, pAb, PM045Y), Anti-Atg5 (pAb, PM050Y), Positive control for anti-LC3 (PM036-PNY).
2.1.2. Cell signaling antibodies (Cell Signaling Technology, 3 Trask Lane, Danvers, MA 01923, USA): Anti-Beclin-1 (pAb, 3738S), LC3A (mAb, 4599S), LC3B (pAb, 2775S).
2.1.3. Abcam antibody (Abcam, Inc., 1 Kendall Square Suite, B2304, Cambridge, MA 02139-1517, USA): p62 (SQSTM1, AB56416).
2.1.4. Sigma antibody (anti-p62/SQSTM1 antibody; P0067)
2.1.5. Secondary antibodies (LI-COR Inc., 4308 Progressive Ave., Lincoln, Nebraska 68504, USA): Goat anti-Mouse (green wavelength) antibody (Li-Cor, C40213-01), Goat anti-Rabbit (green wavelength) antibody (Li-Cor, C30829-02)
2.2. SDS/PAGE and Western Blotting reagents Butanol, 1.5M Tris-HCl (pH 8.8), 1.0M Tris-HCl (pH 6.8), 10% Sodium Dodecyl Sulfate (SDS), N,N,N’,N’-Tetra-methyl ethylenediamine (TEMED, Bio-Rad, 161-0801), 10% ammonium persulfate (APS) (Bio-Rad # 161-0700), 30% acrylamide and bis-acrylamide solution 29:1 (Bio-Rad, 1610156), urea (VWR, BDH4214-500G). Pre-Cast 4-20% gradient gel (Bio-Rad #456-1096 PVDF membrane (Millipore, Immobilon-FL Transfer Membranes IPFL00010) Membrane blocking agent, SeaBlock (ThermoFisher, # 37527); Total protein staining solution (REVERT Total Protein Stain kit, LI-COR Inc., # 926-11010; Pyronin Y (Sigma-Aldrich # P9172)
2.3. Protein concentration determination : Pierce^TM^ bicinchoninic acid (BCA) protein assay kit (Thermo Scientific) (UB276872),
2.4. ELISA commercial kit: For quantifying proteasome (Enzo 20S/26S Proteosome ELISA Kit, Catalog # BML-PW0575-0001)
2.5. *Software:* Image Studio Lite (Ver.4.0) for image analyses. This software is part of Odyssey (LI-COR) image analyzer. Online-based free statistics calculator was used for statistical analysis ( (https://www.danielsoper.com/statcalc/calculator.aspx?id=47)
2.6. *Experimental apparatus:*

2.6.1. Sonic dismembrator (Fisher Scientific, Model: XL2000-350)
2.6.2. Table top centrifuge (Eppendorf, Model: 5418)
2.6.3. Odyssey Infrared Imager (Model : 9120, LI-COR Inc., 4308 Progressive Ave. Lincoln, Nebraska 68504)
2.6.4. Mini Protean III Electrophoresis system (Bio-Rad 165-3301
2.6.5. Electro transfer system (Mini Trans-Blot Electrophoretic Transfer Cell Bio-Rad 170-3930
2.6.6. Multi-well plate reader (Bio-Tek Cytation 5 or Bio-Tek Synergy HT)
3. Procedures:

3.1. *SDS/PAGE and Western Blotting*: The cytosolic proteins of platelet lysates form AD patients and control subjects were separated on a homemade 4-12% gradient gel with 1.5mm width 20 well, MINI Protean II casted gel, using sodium dodecyl sulfate polyacrylamide gel electrophoresis (SDS-PAGE). The apparatus was filled with 1 X electrophoresis buffer. With the gel inside the buffer-containing cell, each lane was loaded with pyronin Y lane marker and 30 μg total protein (1 mg/mL) Electrophoresis was performed at 75 volts for an average of 100 min. until front dye (Pyronin Y) was at the bottom of the gel. The gel was removed from the sandwiched plates and processed for protein transferring to a methanol-activated polyvinylidene difluoride (PVDF) membrane. Electro-transfer process was carried out at 75 volts for 30 minutes. Transfer buffer did not include methanol. The low-temperature of transfer unit was maintained by either inserting an ice-block or placing the transfer unit in an ice-filled container. The PVDF membrane was then stained for total protein visualization by total protein staining kit (REVERT, Total Protein Stain Kit LI-COR Inc., 926-11010). Transferred proteins were imaged by Odyssey (LI-COR) imaging system at the 700 nm set channel. The staining step was optionally removed by incubating with REVERT reversal solution for a maximum of 10 minutes as per protocol provided by the manufacturer. Finally, the membrane rinsed twice with nanopure water. The membrane was blocked in 2 mL of 1:1 SeaBlock /TBS buffer for 1 hour at room temperature (RT) on an orbital shaker. The membrane is directly transferred to a container with 10 mL of 1:1 SeaBlock/TBST and primary antibody dilution in1:500 or 1:1000 while incubating on an orbital shaker for overnight at 4℃. Next day, the membrane was washed with 1X TBST for 20-minutes then incubated horseradish peroxidase conjugated secondary antibody, goat anti-mouse (LC3) and goat anti-rabbit (P62, Beclin-1, and Atg5-12) at 1:10,000 dilution for 1 hour at RT on an orbital shaker. Additional 20-minutes wash with 1x TBST was performed to remove unbound antibodies prior to imaging was completed. The membrane was scanned by Odyssey (LI-COR) imager and analyzed by densitometric quantification using Image Studio Lite (Ver.4.0).
3.2. *26s/20s Proteasome analysis by ELISA method*: Initial proteasome assay conditions were optimized in platelets obtained from Kansas City Community Blood Center (KCCBC). Three sample fractions (i.e., whole platelet lysate, clear supernatant, and membranous pellet) were analyzed by proteasome ELISA kit to identify which of the fraction better represents the proteasome protein profile. 40 μg protein samples/ well were used in ELISA. Assay procedure was performed according to the manufacturer provided protocol. Absorbance was read at 450 nm wavelength using a multi-well plate reader (Bio-Tek, Synergy HT). Standard curve was established and slope of the curve was used to determine the concentration of unknown sample proteasome 26s/20s.
4. *Statistical analyses:* The quantitative analysis for Western blots were performed using Image Studio Lite software program (V 4.0.) The quantitative value of autophagy protein markers represent arbitrary units (a.u.) based on the intensity of the bands. Statistical analysis were performed by a two-tailed unpaired student t-test and Mann-Whitney U test. This was coupled with calculating the value of Cohen’s d and the effect-size correlation, R, using the means and standard deviations of two groups (AD and control). Error bars on all data represents standard error of the mean (±S.E.M)

## RESULTS

We have tested a battery of antibodies for probing autophagy protein purchased from several vendors and compared to each other for selecting the best working antibodies. Individual platelet lysate samples from AD and control group (n=9) were analyzed by Western blotting technique using selected antibodies. The results from three replicates averaged and independently tested as an interval type of data. We only found an increase of LC3-I (p ≤ 0.02) in AD when compared to Control (Fig.2). The select autophagosome target proteins appeared to be elevated; however, there was no statistical difference between AD patients and control group, except LC3-I

**Figure-2.**
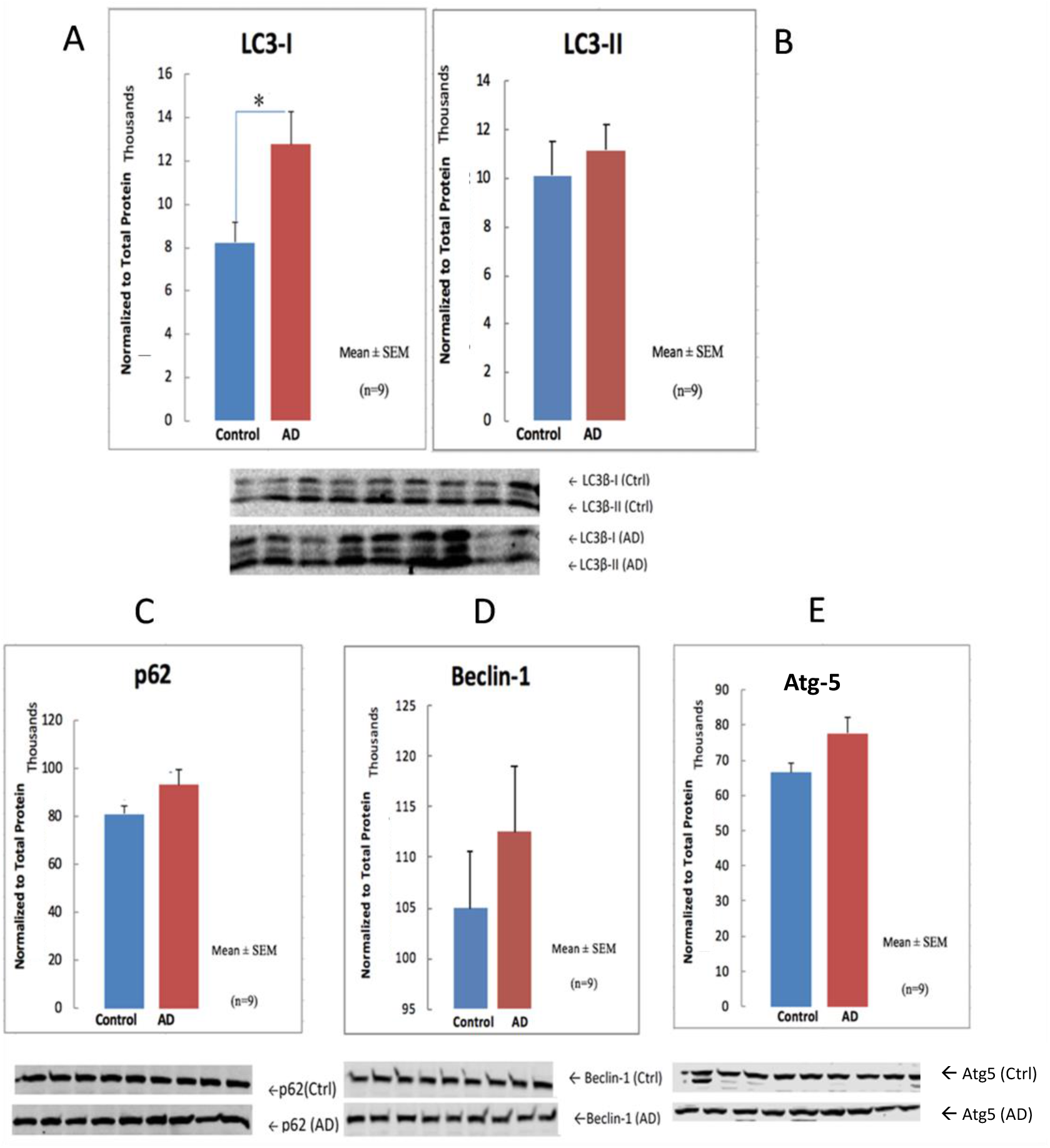
Autophagy target protein profiles in human AD and in non-demented (control) platelet cytosols. The LC3-I, LC3-II, and Atg 5 proteins were probed using antibodies from MBL vendor. The Beclin-1 was detected using Cell-Signaling antibody, and the p62 protein was detected using Sigma-Aldrich antibody. Each samples were analyzed three times and Mean±SEM was presented. The protein band intensities were normalized to total protein staining. Student t-test was employed and only LC3B-1 showed the significance (p≤0.02).

The proteasome concentrations of platelet lysates from AD and age-matched control subjects (n=12) were assessed by ELISA method. Although AD patient platelet lysate proteasome levels were elevated, no statistical difference between patient and control group was obtained (Fig.3)

**Figure-3.**
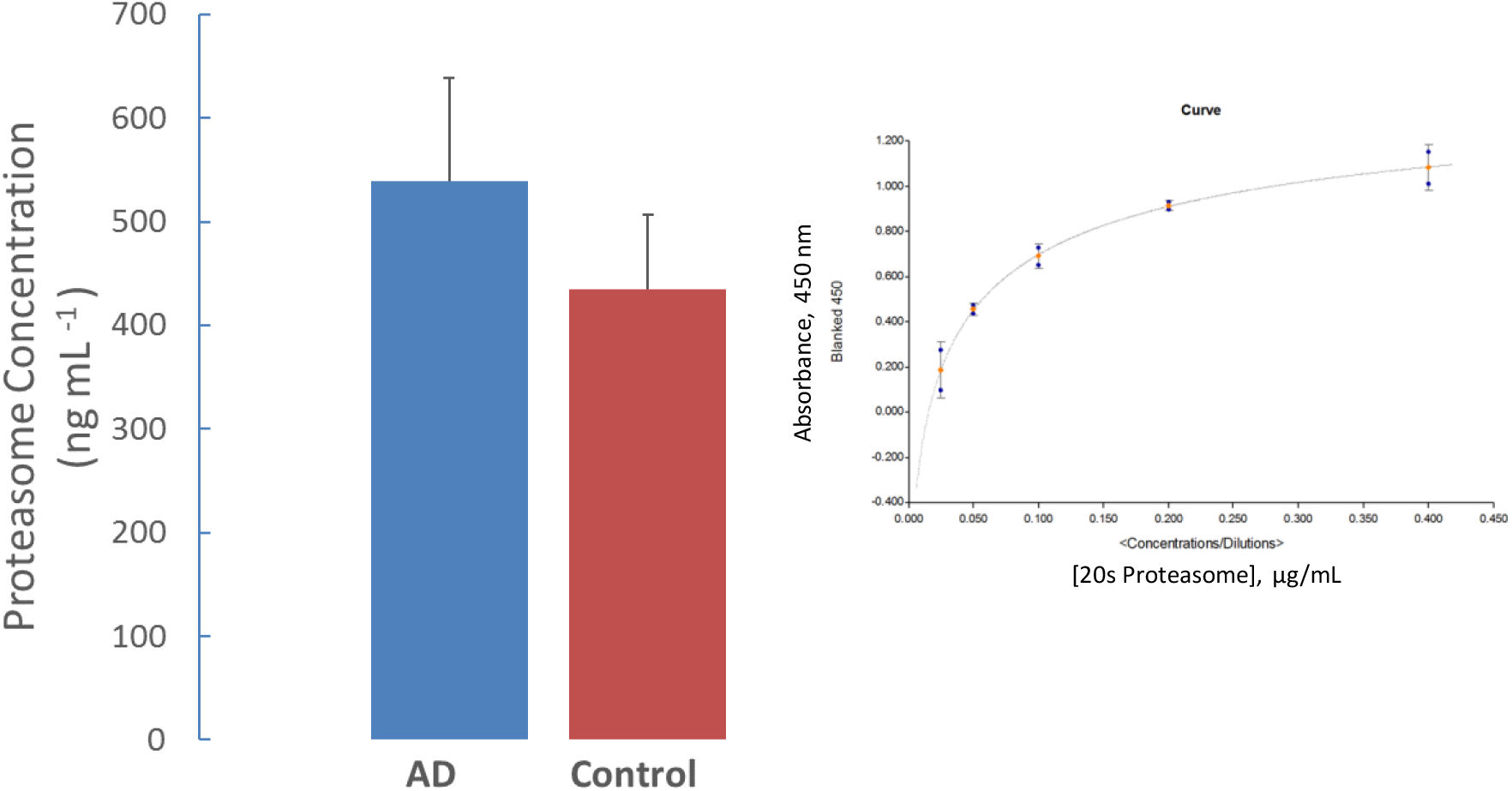
Proteasome concentration measurements in whole platelet lysate. The bar graph represents the proteasome concentration in whole platelet lysate obtained from AD and control subjects (Mean±SEM; n=12). Although an increased proteasome protein profile observed in AD samples, no statistical importance was obtained. The inset figure shows a typical standard curve for quantifying proteasome 20s concentrations.

## DISCUSSION

Aβ_1-42_ is widely understood to accumulate in the early stages of the AD pathology. This malfunctioning protein recruits and triggers microglial, astrocyte facilitated clearance, but soon Aβ_1-42_ depositions overwhelm the response leading to microglia-mediated neuroinflammation ^[14–16]^Aβ_1-42_ also stimulates the catalysis by NADPH oxidase. This enzyme activity increases reactive oxygen species production. The presence of these harmful chemicals activates Atg4, which cleaves and lipidates LC3-I with phosphoethanolamine (PE), as part of the induction of the autophagic pathway. Since AD is a multi-faceted disease, somehow the proteolytic system is suppressed due to downregulation ^[2, 6]^, allowing for the characteristic extracellular amyloid plaques and neurofibrillary tangles to develop.

Some reports also recently discovered that platelets contain a proteolytic system by which hemostasis and thrombosis is acquired ^[17, 18]^. Since we have shown that an elevation of pathological pTDP43 within the platelet cytosol is correlated to hippocampal cortex of AD patients ^[13]^, it was imperative that we analyze these platelet proteolytic systems. We attempted to establish any change of the protein concentrations of proteolytic system, since platelet proteasome concentrations have not been reported prior to our studies. We were unable to assess matching brain tissue proteasome profile from the subjects due to impracticality of obtaining brain biopsy tissue from alive patient and control cohort. In a follow-up study, we consider to include post-mortem brain tissues and matching platelet samples from the same individuals. This approach will provide better connection between brain and platelet proteasome profile. Nevertheless, it was initially believed, hypothesized, based on aforementioned reports, the proteolytic system is somehow altered, and that we would expect to observe some degree of variations in proteasome concentrations.

We employed certain autophagy marker antibodies (i.e. Beclin-1, Atg5, p62, and LC3) to assess the autophagy target proteins in platelets. This was in reference to previous autophagy activity measurements by Gupta et. al, ^[17]^.The antibodies from multiple manufacturers have different affinities for a given sample. A series of antibody dilutions and platelet sample concentrations were optimized in order to match the best antibody /protein combination to be used in the assays. Those results suggested 30 μg whole platelet lysate proteins paired with a primary antibody dilution of 1:1000 was ideal for future analysis.

We evaluated only LC3A and LC3B antibodies in Western blotting. We found that anti-LC3B antibody did not detect a target protein in human platelet lysate samples. Other researchers have stated that out of the three possible isoforms (LC3A, LC3B, and LC3C), LC3A and LC3B autophagosome components exhibit distinct expression patterns in different human tissues ^[19]^. After the final measurement of three replicates, the only notable results found were the proteins levels of LC3-I (a cleaved form of LC3). LC3-II (lipidated LC3-I) is a conjugated protein marker for platelet autophagy system ^[20]^.Since there was no change in levels of LC3-II, the difference in LC3-I between the groups (p ≤ 0.021) could be a result of autophagic pathway being blocked in its initial steps in AD ^[21]^. The overarching implication we believe is that in AD, probably to alleviate aggregate stress, platelet allocated autophagy may be functionally upregulated and induced but inhibited in early or late steps of the pathway^[20]^. Our results seem to be inconclusive at this stage based on exclusively LC3 immunoblotting results. The immunoreactivity of LC3-I and LC3-II are different. LC3-II tends to be more sensitive to antibodies^[21]^. A stimulated autophagic pathway is represented by reduction of autophagy marker LC3II. So, when we analyzed that there was no change in LC3-II, its more indicative of no detection of flux than the changes of LC3-I. Therefore, an autophagy activity analysis assessment should be performed in freshly isolated intact platelets in a serious of follow-up study.

We assessed the proteasome concentration in platelet lysates. Poteasome 20S protein levels were measured by enzyme-linked immunosorbent assay (ELISA) techniques in cell lysates, supernatants, and pellets in KCCBC obtained healthy samples in initial phase of these studies. We relied on supernatant isolations of platelet lysates to measure AD profiled proteasome concentrations. We found no statistical difference (p=0.373) between AD patients and age-matched control subjects in these platelet samples based on a two-tailed, student t-test and Mann-Whitney U test.

For effectively assessing the relationship of platelet proteolytic machinery protein levels in AD rather than the use of only “p” values, we incorporated effect sizes in our data analysis ^[22, 23]^. In addition to selecting a student t-test (interval, parametric), we also employed the Mann-Whitney (ordinal, non-parametric) level of measurement. This was justified by the expression of a large variation in protein concentrations between individual samples within each group. This means our sample did not meet the parametric assumption of equal variances. This adjustment ascribed the data into ranked categories before comparing the median values between the groups.

Despite the statistical insignificance between the two group’s means of Atg5, p62, Beclin-1, a measure of strength of an effect points may be more meaningful. A moderate to large standard deviations (d = 0.41–1.0) and correlations (R = 0.36-0.44) between AD and control were detected, denoting a probability that there is a difference in autophagic pathway protein levels. Our sample size was not large enough to produce statistical significance. On the other hand, the insignificance found between AD and control regarding LC3-II protein bands was coincided with a small magnitude of effect between groups. (d = 0.27, R = 0.13, p = 0.58) (Fig.1B) In proteasome concentration profile assay, we have observed a statistically insignificance with a small group effects (p=0.373, d = 0.34, R = 0.17).

In this study, there are some caveats and setbacks, rendering our results with a low difference. First, our continuous and averaged data comparison was statistically insignificant for all except one based on our sample cohort (n=9 for autophagosome and n=12 proteasome analyses). As mentioned before, human biological samples do have a substantial variation in concentration and effect between each sample and group measurements. Especially with a small effect size for the main autophagy monitor, LC3-II, increasing the n value to ~220 per group (n=220) should produce a significant t-test at a probability level of 0.05 for all samples ^[24]^. Secondly, as it has been reported that LC3-I is unstable and sensitive to freezing-thawing cycles or SDS buffered cocktail ^[25]^. Future samples should be freshly assessed and not exposed to a repeated freeze-thaw cycle. Thirdly, immunoblotting has its limitation of determining autophagic fluctuations. As we understand that LC3-II is the go-to marker, an increase in concentration could be due to correlate with autophagosome accumulation but autophagic flux is not guaranteed ^[26]^. In more relevance to AD neuropathology, beclin-1 is downregulated in this disease; however, LC3-II can still form without a phagophore on ectopic membrane fragments ^[21, 25]^. Beclin-1, an adaptor protein via its partner proteins, can either stimulate or suppress the onset of autophagy ^[27]^. In this study, we observed that beclin-1levels were elevated in AD patient’s platelet. We are unable to offer an explanation on beclin-1 behavior, because we don’t know the autophagy activity profile at this stage.

It is known that a neurodegenerative condition such as AD effects the periphery by increase in thrombin and von Willebrand factor (vWF) protein. Thrombin and vWF are platelet activators ^[18]^. In AD, the patients are known to express an increased amount of vWF, which is partially used to convert quiescent platelets into activated platelets ^[28]^. Hence, activated platelets of these patients produce and release more Aβ into circulation compared to the controls ^[29]^. Aβ usually interacts with fibrin thereby promoting coagulation and fibrin aggregation. Therefore, an increase of Aβ could mean an increase in platelet aggregation that implies an alternate flux in hemostasis. We did not analyze Aβ peptide levels in platelet lysates of AD patients and age-matched control subjects in this study; however, we plan to include the platelet lysate Aβ_1-42_ measurements in a follow-up study.

Since our results are supposed to be a testament of basal autophagy in quiescent platelets, AD derived samples might have already been activated when obtained from the patients. Activated platelets have increased autophagic pathway activity ^[18]^. Therefore, the significant measurements might be due to that previous condition. One of the way to normalize this variable would be to activate the control platelets before any protein concentration measurements. An alternative approach would be to inhibit the platelet activation factor by including an inhibitor (PGI_2_) during the platelet isolation from whole blood. Platelet proteolytic system analysis has allowed us to differentiate probable proteostasis between AD and control in cross-sectional fashion. One of the possibilities that can be discerned from our study is that there may be elevated induction of autophagy, beyond basal amount.

In light of this information, platelet autophagic profile may be similar to neurons of this disease profile. This study was meant to extrapolate the extent to which AD can influence the state of these systems during an elevated presence of disease-related proteins. Autophagy flux and proteasome assessment in intact platelet are necessary for obtaining more evidences that are conclusive. A similar cross-sectional study with a few adjustments should be considered before testing other neurodegenerative disease group in a similar fashion.

## DECLARATIONS

### Authors’ contributions

Muriu, RG made substantial contributions to conception and design of the study and performed data analysis and interpretation

Sage, JM performed data acquisition, as well as provided administrative, technical, and material support Agbas, A conceptualized the study, analyzed the data, and wrote the manuscript

### Availability of data and materials

Not applicable

### Ethical approval and consent to participate

The authors state that they have obtained appropriate institutional review board approval (KUADC#11132) or have followed the principles outlined in the Declaration of Helsinki for all human or animal experimental investigations. In addition, for investigations involving human subjects, informed consent has been obtained from the participants involved.

### Consent for publication

Not applicable

### Copyright

© The Author(s) 2019

